# Blocking uncertain mispriming errors of PCR

**DOI:** 10.1101/2024.04.19.590219

**Authors:** Takumi Takahashi, Hiroyuki Aoyanagi, Simone Pigolotti, Shoichi Toyabe

## Abstract

The polymerase chain reaction (PCR) plays a central role in genetic engineering and is routinely used in various applications, from biological and medical research to the diagnosis of viral infections. PCR is an extremely sensitive method for detecting target DNA sequences, but it is substantially error-prone. In particular, the mishybridization of primers to contaminating sequences can result in false positives for virus tests. The blocker method, also called the clamping method, has been developed to suppress mishybridization errors. However, its application is limited by the requirement that the contaminating template sequence must be known in advance. Here, we demonstrate that a mixture of multiple blocker sequences effectively suppresses the amplification of contaminating sequences even in the presence of uncertainty. The blocking effect was characterized by a simple model validated by experiments. Furthermore, the modeling allowed us to minimize the errors by optimizing the blocker concentrations. The results highlighted an inherent robustness of the blocker method, in that fine-tuning of the blocker concentrations is not necessary. Our method extends the applicability of PCR and other hybridization-based techniques, including genome editing, RNA interference, and DNA nanotechnology, by improving their fidelity.

**Significance:** The applications of PCR are increasing day by day, and there is a need to suppress PCR errors to improve the accuracy of PCR-based techniques and broaden their applicability. The blocker method has been developed to substantially suppress mispriming. However, the method requires prior knowledge of the contaminating sequence, which limits its applicability. We successfully demonstrate that adding a combination of multiple blocker sequences can substantially suppress PCR errors, even when we have only partial information about the contaminating sequences. We also construct a biophysical model of the blocking effect, which allows us to find the optimal blocker combinations that minimize the PCR error. Since the method targets hybridization, it is readily applicable to a wide range of biotechnologies.

## 1 Introduction

Polymerase chain reaction (PCR) is a technique for amplifying DNA strands characterized by specific sequences^1^. In PCR, a pair of short strands, called primers, are mixed in the reaction solution. They specify the sequence region to be replicated and trigger DNA replication. By running several replication cycles, PCR is able to exponentially amplify a DNA sample. This exponential amplification makes PCR extremely sensitive, but also error-prone. On the one hand, the incorporation of incorrect nucleotides during polymerase extension can be significantly suppressed by using a high-fidelity polymerase and carefully tuning the reaction conditions. On the other hand, errors can be caused by mishybridization of primers onto contaminating strands^2^ (Fig.1a). In practical applications, test samples are often far from being pure, and contain a variety of DNA sequences. Primers readily hybridize to contaminating sequences similar to the target sequence, resulting in false positives in viral testing.

**Figure 1.**
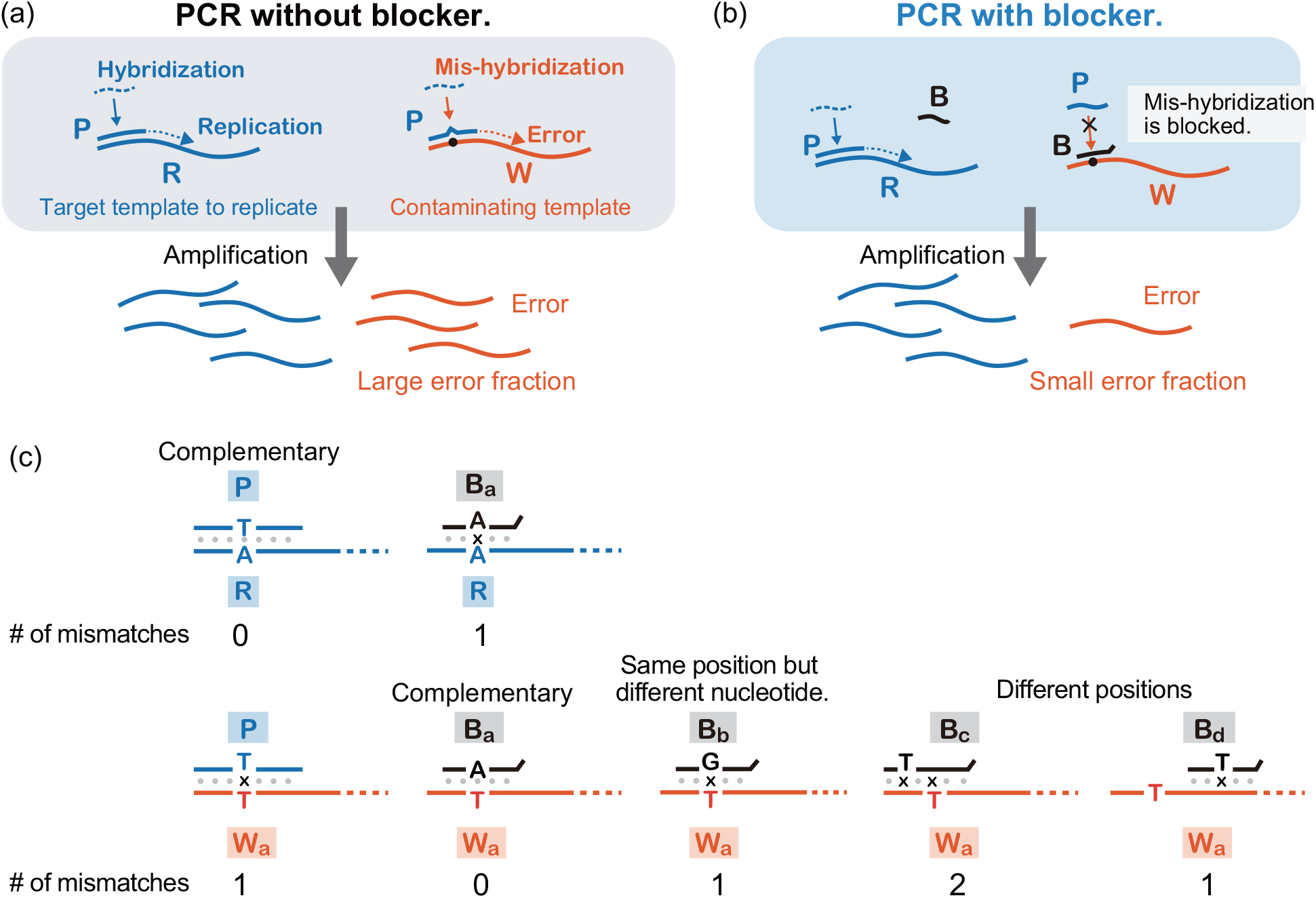
PCR and blocker method. (a) Mishybridization of the primer strands (P) to the contaminating template strands (W) results in errors. (b) Blocker method. The sequence of the blocker (B) is designed to be complementary to the sequence of W and blocks the primer binding to W. Two-base floating end is attached at the 3’-end of B to prevent priming at this end. (c) Different sequences W_*i*_ and B _*j*_ are labeled so that *i* = *j* when W_*i*_ and B _*j*_ are complementary. The number of mismatches changes depending on the relative positions of the mutation in W and the target mutation in B.

The blocker or clamping method was developed to suppress mispriming in PCR^3–6^. (Fig.1b). In this method, a nucleic acid sequence called a blocker is added to the PCR mixture. The blocker sequence is designed to be complementary to the contaminating sequence. It suppresses errors by blocking mishybridization of the primer to the contaminating sequence (Fig. 1c). A chimeric strand of DNA and locked nucleic acids (LNA)^7^ or DNA and peptide nucleic acids (PNA)^8^ are often used as blockers due to their high specificity^3,9,10^. In our previous work, we established that blocking combines energetic and kinetic discrimination toward correct hybridization to the target sequence^10^. Because of this mechanism, error suppression remains effective even at relatively low temperatures, and therefore, it is expected to be applicable to thermally sensitive systems such as biological cells. Since the blocker sequence is designed to be complementary to the contaminating sequence, the blocker method requires to know the contaminating sequence in advance. This limitation can be quite severe in realistic situations where there are multiple possible contaminating sequences, but we do not know which sequences are contaminating each sample.

This work extends the blocker method to situations where the information about contaminating sequences is limited. For this purpose, we demonstrate the effectiveness of the blocker combinations in suppressing errors. At the same time, we construct a simple biophysical model and validate the model through extensive experiments to characterize the error suppression by blocker combinations. Finally, we attempt to minimize the errors by optimizing the blocker concentrations.

We specifically consider the following scenario. A PCR mixture contains a target sequence to be detected, R, and a contaminating sequence, W_*i*_, randomly sampled from *N* possible sequences with probability *p*_*i*_ (1≤ *i* ≤*N*). The primer P is complementary to the primer-binding region of R. The contaminating sequence has a corresponding sequence region with single or multiple mutations, which cause mismatches in the hybridization with P. The blockers are labeled as B_*i*_, such that B_*i*_ is perfectly complementary to W_*i*_ (Fig. 1c).

## 2 Model

Our method is based on adding a combination of blocker sequences {B_*i*_ } that are complementary to all the possible contaminating sequences {W_*i*_}. Throughout this paper, we use to {·}denote a set. The primer we use in this work is 20 bases long, which is typical for PCR virus tests. Therefore, there exist in principle 3^20^ possible W sequences. However, the binding strength of P to W with two or more mismatches is negligible in standard PCR assays. Therefore, in practice, we only need to worry about the contaminating sequences with a single mismatch with respect to R, which are limited to 3× 20 = 60 sequences.

We define a growth rate *r* as the number of strands copied in a PCR cycle divided by the number of template strands. With this definition, perfect PCR efficiency corresponds to *r* = 2. We call *r*_R_ and *r* 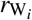 the growth rates of R and W_*i*_, respectively. Then, the error fraction is defined as^11^

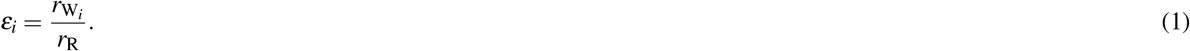

Once mispriming occurs, the product works as a normal template in successive cycles. That is, the product contains the correct primer-binding sequence. Therefore, the mispriming typically happens only for the contaminating sequences and not for the PCR products. That is, the overall error of the PCR assay should show not a power-law-like but linear dependence on *ε*_*i*_.

We call *b*_*i*_ the concentration of B_*i*_ and *ε*_*i*_({*b*_*k*_}) the error fraction associated with W_*i*_ in the presence of a blocker distribution {*b*_*k*_}. We evaluate the error by either the mean error fraction or the maximum error fraction, defined as

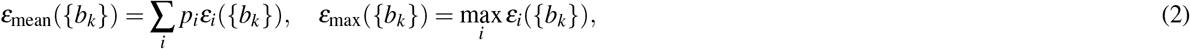

respectively, under the constraint that the total concentration *b*_tot_ = ∑_*i*_ *b*_*i*_ is fixed. This constraint reflects the limitation that too many blocker strands also suppress the replication of R, as we will see (Fig. 2a). The choice between focusing on *ε*_mean_ or *ε*_max_ depends on the purpose of the PCR assay. One may want to minimize *ε*_max_ in diagnosis to avoid false positives. For DNA cloning by PCR, reducing *ε*_mean_ may increase the product amount of the right sequence more than reducing *ε*_max_, although this may depend on the system.

**Figure 2.**
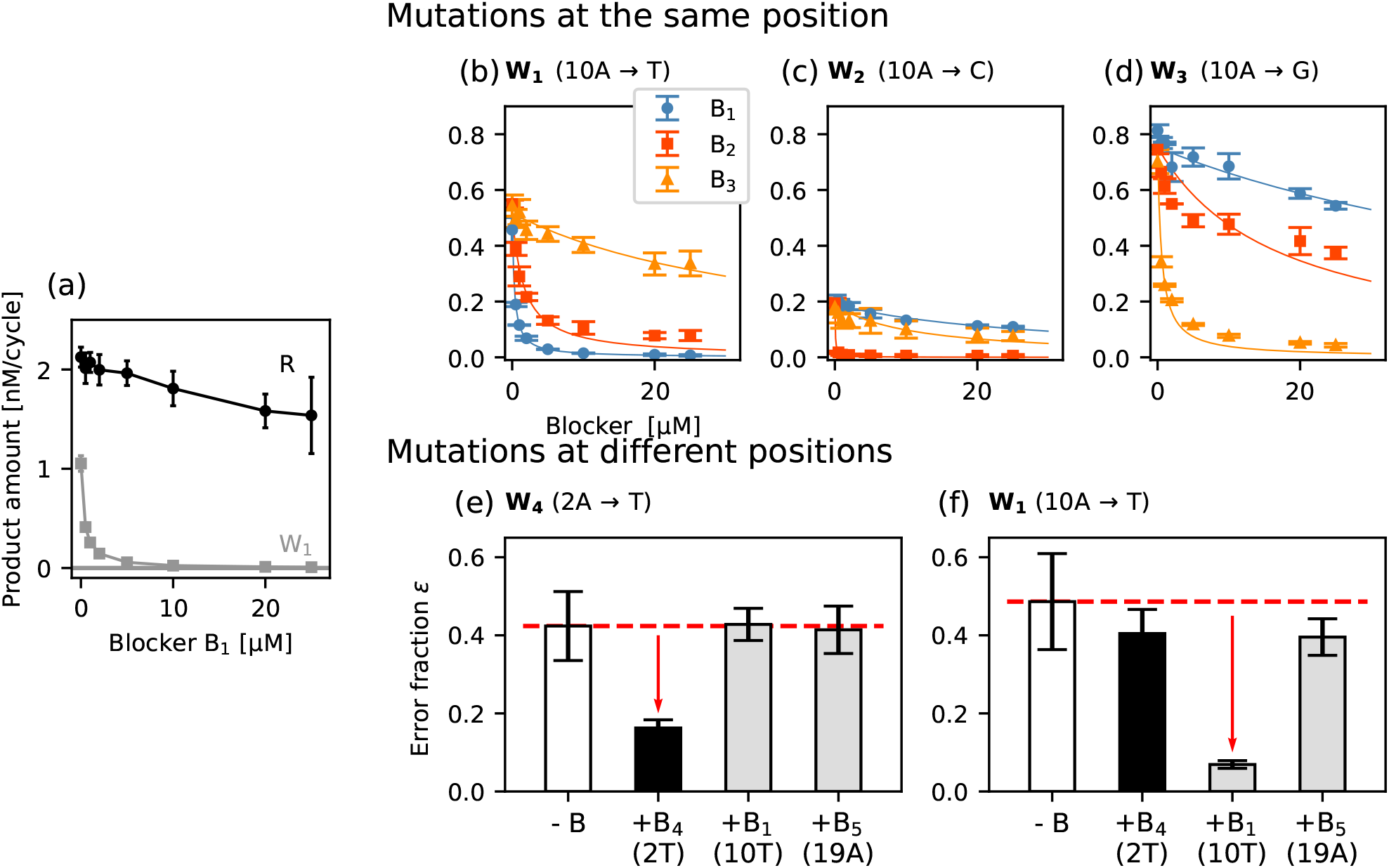
Error suppression by a single blocker sequence. **a**, The production amounts of R and W_1_ per cycle in the presence of blocker sequence B_1_. **b**–**d**, Dependence of the error fractions *ε* on blocker concentration for different combinations of W and B, where the target mutations of B are all at the same position in W. The subscript *i* of W_*i*_ is used for labeling different W sequences. B_*i*_ is the blocker sequence complementary to W_*i*_. The expression, for example, W_1_ (10A→T) indicates that A, the tenth base from the 3’ end in R, is mutated to T in *W*_1_. The solid curves are fitting curves by Eq. (4) with the fitting parameter *K*_*i,j*_. **e, f**, Error suppression including the situations where the positions of the target mutations of B are different from the mutated position of W. The concentration of the blocker is 2 µM. Error bars indicate standard deviations. Each data point represents an average over three independent experiments.

We consider a biophysical model for evaluating *ε*_*i*_({*b*_*k*_}). First, we consider the situation with a single B sequence and then extend the model to multiple B sequences. In experiments, the blocker concentration is relatively high (∼µM). We therefore assume that the blocker binding and dissociation are always equilibrated^10^. Then, the growth rates are given by

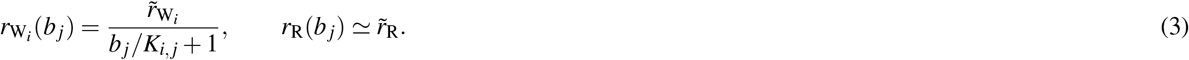

Here, *K*_*i,j*_ is the dissociation constant associated with the binding of W_*i*_ and B _*j*_. The factor 1*/*(*b*_*j*_*/K*_*i,j*_ + 1) represents the fraction of the free W_*i*_ that is not bound by B _*j*_. We assumed that *b* _*j*_ is not extremely large. 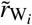 is the growth rate in the absence of the blocker sequences. We assume that the primer then binds to free W_*i*_, initiating the polymerase extension. Because the binding of B _*j*_ to R is negligible under the present condition (Fig. 2a), the replication of R_*i*_ is not blocked by the blockers;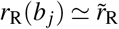

Combining Eqs. (1) and (3), we obtain an expression for the error fraction in the presence of a blocker B _*j*_:

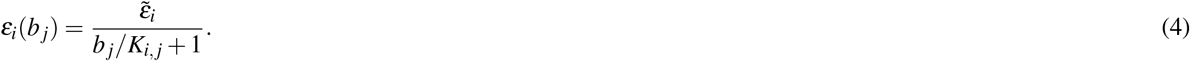

Here, 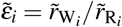 is the error fraction in the absence of blocker sequences. Thus, the model predicts that the blocking strength of W_*i*_ by B _*j*_ is essentially determined by *K*_*i,j*_. In the presence of multiple blockers, Eq. (4) becomes

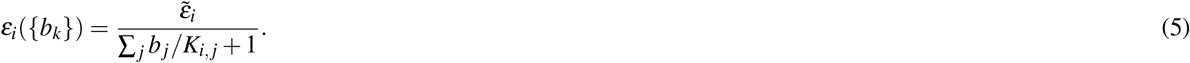

Since *b* _*j*_ and *K*_*i,j*_ are non-negative, Eq. (5) implies 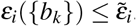. As expected, the error is always reduced by the addition of blockers.

## 3 Experiments

We performed experiments to characterize the effect of the blockers on the error fraction; to validate Eqs. (4) and (5); and also to evaluate 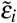 and *K*_*i,j*_. We performed a linear PCR using a single primer instead of a primer pair since the error fraction is directly evaluated from the product amounts in the linear PCR (Fig. S1). The reaction mixture contains 2.5 nM R template, 2.5 nM W_*i*_ template, 100 nM primer, thermostable (Taq) DNA polymerase enzyme, and standard buffer mixture. Blocker sequences are added depending on the experiments. See the Methods and SI for details.

We use W sequences with mutations at the positions where the original nucleotide of R is either A or T. Mutations at A and T cause more frequent errors than those at C and G positions due to their lower binding stability. The reason for this choice is to focus on the worst cases for practical assays.

### 3.1 Dependence of PCR error on blocker concentration

We measured the amount of the products in the presence of a blocker sequence (Fig. 2a) to evaluate the dependence of the error fractions on the blocker concentrations for different combinations of W_*i*_ and B _*j*_ (Fig. 2b–d). The error fraction *ε*_*i*_(*b*_*j*_) decreased significantly with *b* _*j*_ for the complementary pair *i* = *j*. This dependence is well fitted by Eq. (4), supporting the validity of the model.

We distinguish two cases for non-complementary pairs of W_*i*_ and B _*j*_ (*i* ≠*j*). If the mismatch positions of *i* and *j* are the same, the blocker still suppresses the error (Fig. 2b–d). This is because the binding of B _*j*_ to W_*i*_ is not negligible in this case. In particular, purine-pyrimidine pairs (A and C) and (G and T) bind weakly even though they are not complementary. This crosstalk effect between B and W explains the tendency of the error fractions in Fig. 2b–d. For example, in Fig. 2b, B_2_ suppresses errors more than B_3_. This tendency is due to the stronger binding of B_2_, which has G at the corresponding mutation position, to W_1_ than B_3_, which has C. The error fraction without blocker is the highest in Fig. 2d because G, the mutation in W_3_, and T of the primer at the corresponding position bind to some extent. Equation (4) fits the results well for such pairs.

When the mismatch positions of *i* and *j* are different, error suppression is negligible (Fig. 2e, f). This is because the number of mismatched nucleotides is two (see, for example, W_a_ and B_c_ in Fig. 1c) and their hybridization is unstable. This may not be the case if the two mismatches are far apart (see W_a_ and B_d_ in Fig. 1c). In such cases, however, the primer binding region is not sufficiently covered by the blocker, and blocking is again not effective.

When the blocker concentration is higher than 10 µM, the replication of R is also suppressed (Fig. 2a). Therefore, we used *b*_tot_ = 10 µM in the following experiments.

### 3.2 Estimation of dissociation constants *K*_*i,j*_ from sequences

Directly measuring *K*_*i,j*_ for all the 60×60 pairs of W_*i*_ and B _*j*_ is unpractical. Based on the results in Fig. 2e, f, we set *K*_*i,j*_ = ∞for non-complementary pairs with different mismatch positions. For other pairs, we estimated the *K*_*i,j*_ values by the nearest neighbor method^9,12,13^ with LNA parameters including mismatches (Fig. 3). We found that the values estimated by the nearest neighbor method are similar to those obtained by the fitting of Eq. (4) in Fig. 2b–d.

**Figure 3.**
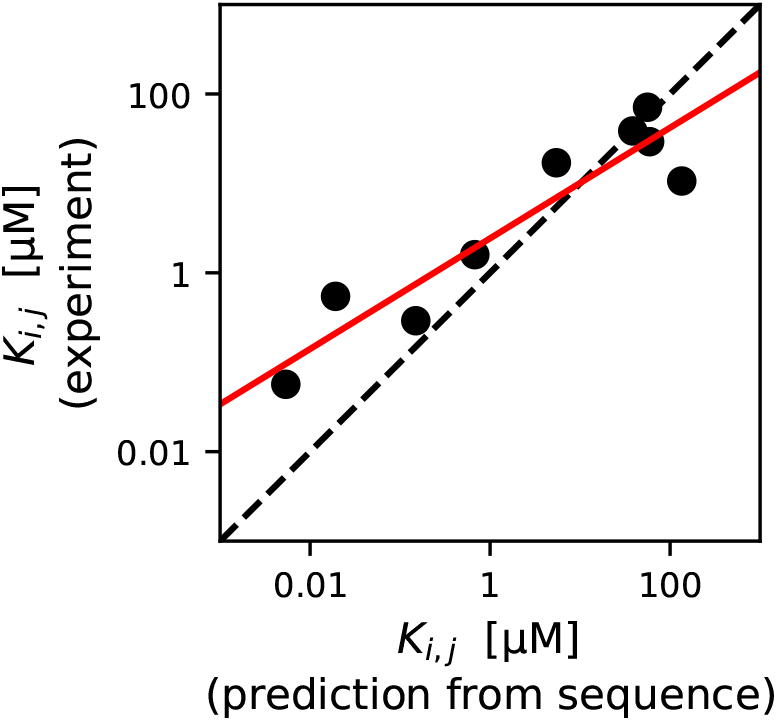
Experimental vs. predicted values of the dissociation constants *K*_*i,j*_. The experimental values *K*_*i,j*_ are obtained by fitting Eq. (4) to the experimental curves in Fig. 2b–d. The predicted values are the estimation from the sequences by the nearest-neighbor method^9,12,13^. We used the BioPython package with modifications^14^. The dashed line indicates the equality between the experimental values and predicted values. The solid line is a fit by a curve linear in the log scale: ln *y* = *a* ln *x* + *b* with *a* = 0.619 and *b* = −4.38. Here, *x* and *y* are in the units of M.

Broadly, this result supports the validity of the model given by Eq. (4) for the blocking effect. However, there are small deviations between the experiments and predictions. The deviations could be due to the fact that the parameters of the nearest neighbor method are not optimized for our experimental conditions. For better estimation, we used a linear fitting in the log scale (red solid curve in Fig. 3) to calibrate the *K*_*i,j*_ values estimated by the nearest-neighbor method.

### 3.3 Estimation of error fraction without blocker 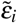

We measured the dependence of 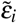 on the position and nucleotide species of the mismatch of W (Fig. 4). We observed a tendency for 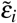 to be small at both the 5’ and 3’ ends of the primer binding region of W. This is because a mismatch near the ends causes the ends to flap and reduces stability more than internal mismatches. In particular, rigid hybridization at the 5’-end of the primer binding region of W (3’-end of the primer) is necessary for the polymerase to initiate extension^15–17^.

**Figure 4.**
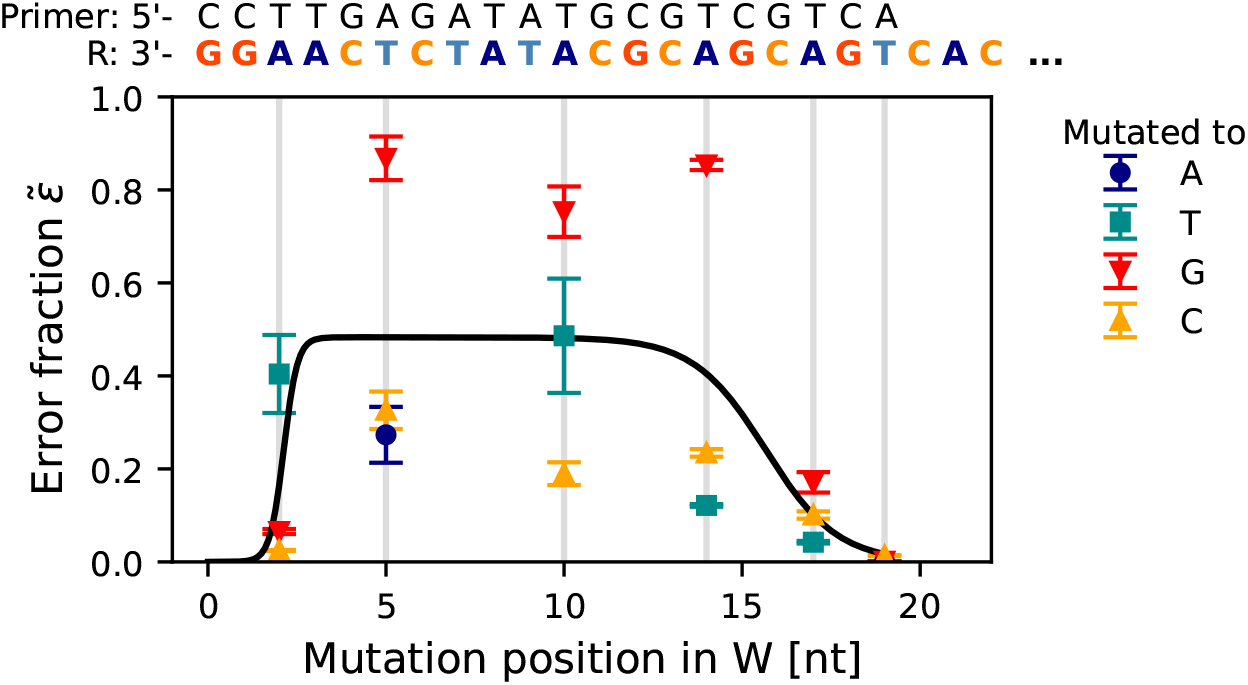
Dependence of error fraction 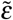 on the mutation position and the nucleotide species of W in the absence of the blockers. Solid line is a fitting curve *f* (*x*) = *p*[1 + tanh(*q*(*x*− *r*))][1− tanh(*s*(*x*− *u*))] with the fitting parameters *p* = 0.12, *q* = 2.6, *r* = 2.1, *s* = 0.50, and *u* = 16. We chose this function since it can reasonably model the observation that the error fraction is small at the edge and similar in the bulk sequences. Each symbol corresponds to a different nucleotide type in the W sequence. Error bars indicate standard deviations. The number of independent experiments is three for each point. The sequences of R and P are shown on the top for the reference.

The value of the parameter 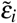 depends on the nucleotide species (Fig. 4). However, this dependence is hard to model accurately. For example, the sequence near the mismatch position should affect 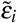.Measuring all sixty values of 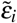 is not practical. We could calculate *K*_*i,j*_ from the sequences because the blocker concentration is sufficiently large, and the equilibrium is expected for the blocker binding. However, equilibrium is not expected for primer binding because the primer concentration is not large enough^10^. Therefore, it is not straightforward to estimate 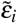 only from the sequences. As a reference, the free energy difference ΔΔ*G* between the hybridization P:R and P:W_*i*_ are calculated by the nearest neighbor method^12,13^ (Fig. S3). The results show a similar tendency as 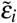 in Fig. 4. This implies that 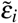 depends largely on the hybridization stability.

Here, we assume a simple model where 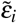 depends only on the mismatch position and neglect the dependence on the nucleotide species. To account for the flat dependence on position in the internal region, we fit 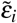 with a combination of two hyperbolic tangent functions (Fig. 4). We, however, observed a significant variation around the fitted line. We will see how the variation affects the optimization of the blocker concentrations.

### 3.4 Suppression of errors with a specific error by blocker combination

In order to test the effectiveness of the blocker combinations and the model (Eq. (5)), we contaminated one of three specific W sequences and evaluated error suppression by a combination of three blocker sequences targeting the three W sequences (Fig. 5a–c). The sum of blocker concentrations was kept at ∑ _*j*_ *b* _*j*_ = 10 µM. We only tested the W sequences with the same mutation positions since blocking is ineffective if the target position of B differs from the mutation position of W, see Fig. 2e, f. We found a gradient of the error fraction in a ternary diagram. The steepest gradient is obtained along the axis of the blocker complementary to the contaminating sequence (*i* = *j*), as expected.

**Figure 5.**
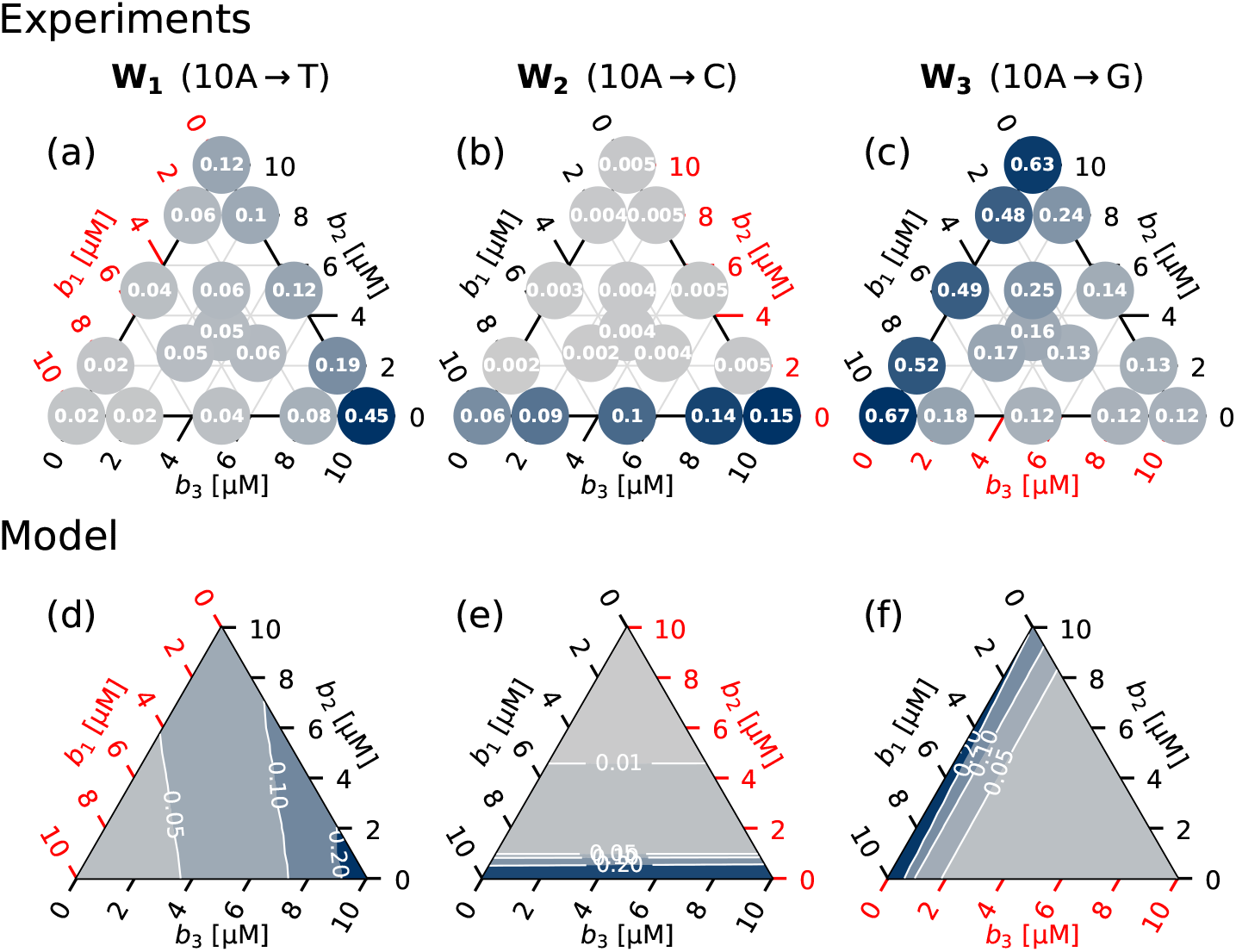
Error suppression by combinations of multiple blocker sequences. Experimental data (top) and corresponding model calculation by Eq. (5) (bottom). The error fractions without blockers 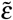 are 0.52 (a), 0.19 (b), 0.75 (c), and 0.48 (d–f). The numbers in the circles in (a–c) are the mean error fractions of three independent experiments. The blockers highlighted by red color are complementary to the contaminating sequences. *b*_tot_ = 10 µM. The plots are colored according to the error fractions normalized by the maximum error fraction in each plot.

We now evaluate *ε*_*i*_({*b* } _*j*_) by Eq. (5) using *K*_*i,j*_ calculated by the nearest neighbor method with calibration (Fig. 3) and 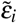 modeled in Fig. 4. We found that Eq. (5) qualitatively reproduces the experimental results under different conditions (Fig. 5d–f).

The absolute values show some discrepancy between the experimental results and the model calculations. The main source of the discrepancy seems to be the approximation by neglecting the dependence of 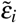 on the nucleotide species (Fig. 4).

### 3.5 Suppression of errors with partial information by blocker combination

We test if the blocker combinations are still effective when the information about the contaminating sequences is limited. We apply this approach to three situations with different mismatch patterns (Fig. 6), where we have partial prior knowledge of the errors. In both examples, we considered the worst case, where the contamination probability is uniform, *p*_*i*_ = 1*/N*. The results are summarized in Table 1. In each scenario, we calculated *ε*_mean_ and *ε*_max_ from experimentally measured *ε*_*i*_ using Eq. (2).

**Table 1.**
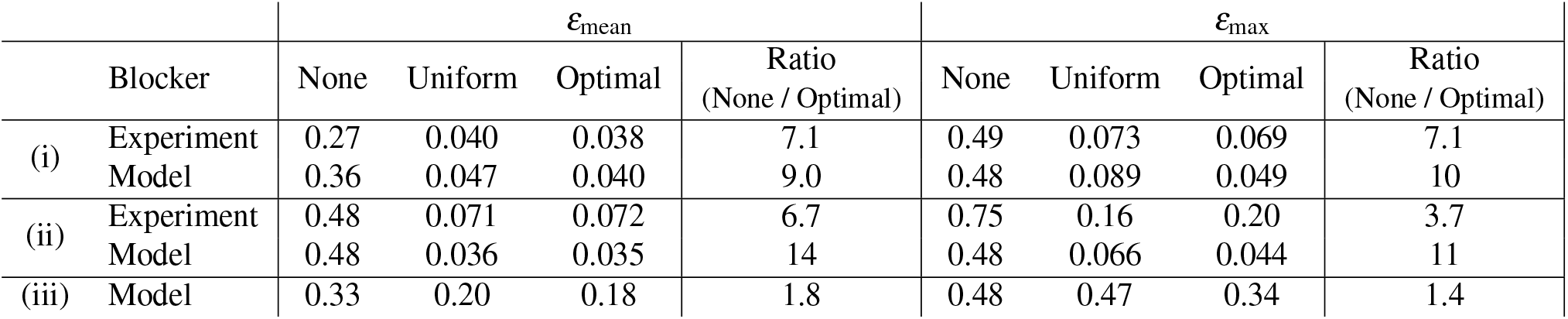
Summary of error suppression. *ε*_mean_ and *ε*_max_ without blocker (None), with blocker combinations with the uniform concentrations (Uniform, *b*_*k*_ = *b*_tot_*/N*), or with the optimal concentrations (Optimal,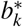). (i) The mutation is located in one of the three positions. The nucleotide species is specified (Fig. 6a, b). (ii) The mutation is located in the specified position, but the nucleotide species is not identified (Fig. 6c, d). (iii) Neither position nor nucleotide species are specified. The experimental values are obtained by interpolating the blocker concentrations in Fig. 6.

**Figure 6.**
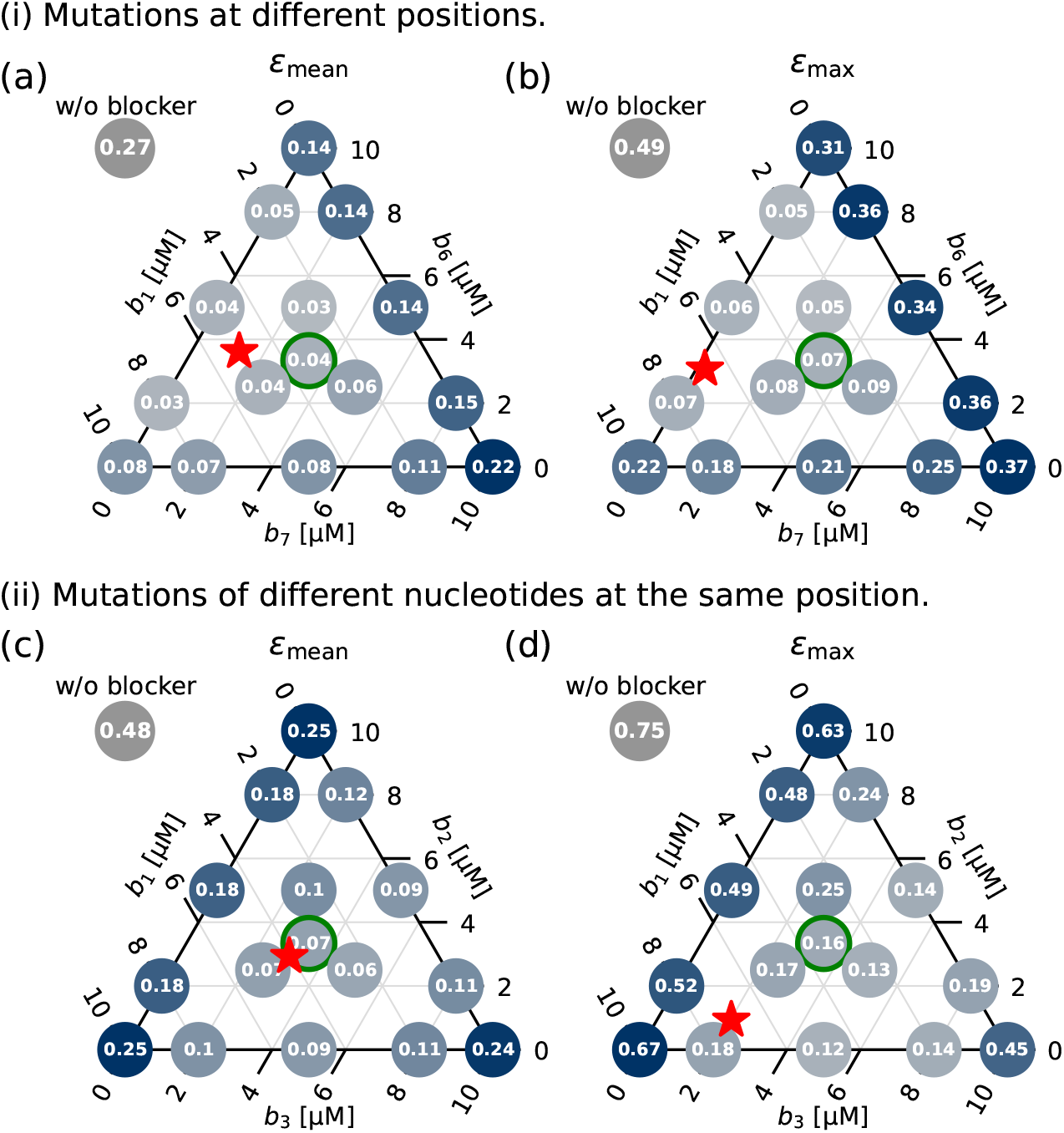
*ε*_mean_ and *ε*_max_ values in the presence of blocker combinations obtained by experiments. Mutations are located at different positions (a, b) or same position with different nucleotides (c, d). The blockers *b*_*i*_ are complementary to W_*i*_. W_1_: 10A →T, W_2_: 10A→ C, W_3_: 10A→ G, W_6_: 5T→ A, W_7_: 17A →T. We measured *ε*_*i*_({*b*_*k*_}) and plot *ε*_mean_ = ∑_*i*_ *ε*_*i*_({*b*_*k*_ })*/*3 and *ε*_max_ = max_*i*_ *ε*_*i*_({*b*_*k*_ }). The circles indicated by green corresponding to the uniform blocker concentrations. The stars indicate the optimal blocker concentrations calculated by the model. The corresponding error fractions are 0.038 (a), 0.069 (b), 0.072 (c), and 0.20 (d), which are calculated by a linear interpolation. See Figs. 7a, b, S4a, b for corresponding model calculations.

#### (i) The mutation is located in one of the three positions. The nucleotide species is specified (Fig. 6a, b)

In this scenario, B _*j*_ has a negligible error suppression effect on W_*i*_ that does not match (*i* ≠*j*) as seen in Fig. 2e, f. Hence, the effect of the blockers can be decomposed as *ε*_*i*_({ *b*_*k*_}) = *ε*_*i*_(*b*_*i*_), leading to a simple superposition *ε*_mean_ = ∑_*i*_ *p*_*i*_*ε*_*i*_(*b*_*i*_).

Taking a uniform blocker concentration *b*_*i*_ = *b*_tot_*/N* as a reference (indicated by green circles in Fig. 6), we have a significant error reduction with the reduction ratio of 6.8 for *ε*_mean_ and 6.7 for *ε*_max_ (Table 1). This significant error suppression shows the effectiveness of our approach.

#### (ii) The mutation is located in the specified position, but the nucleotide species is not identified (Fig. 6c, d)

As shown in Fig. 2b–d, *B*_*j*_ can suppress the replication of *W*_*i*_ even if *i* ≠ *j* when the mutations of *i* and *j* are located at the same position. Hence, the error suppression is more complicated than in case (i). With uniform blocker concentrations, we obtained significant reduction ratios of 6.8 for *ε*_mean_ and 4.7 for *ε*_max_ (Table 1).

Thus, in these distinctive scenarios with partial information about the contaminating sequences, the addition of blocker combinations significantly reduced errors, supporting the practical effectiveness of the method.

We calculated the error pattern by the model. By substituting Eq. (5) to Eq. (2), we obtain

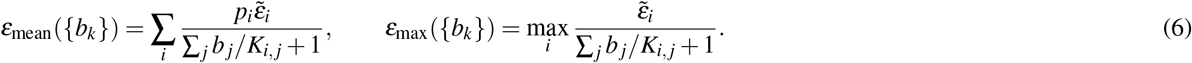

The parameters 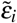 and *K*_*ij*_ could be calculated by the model in Fig. 3 and 4, respectively, for the given primer sequence.

Figures 7a, b and S4a, b show the model calculations corresponding to Fig. 6a, c, b, and d, respectively. The model qualitatively reproduced the error pattern obtained by experiments. We observed a quantitative discrepancy for *ε*_max_. This is possibly due to neglecting the nucleotide dependence of 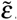. In this particular scenario, this assumption could significantly affect the estimation. The estimation of the parameters can be improved for each system if necessary.

**Figure 7.**
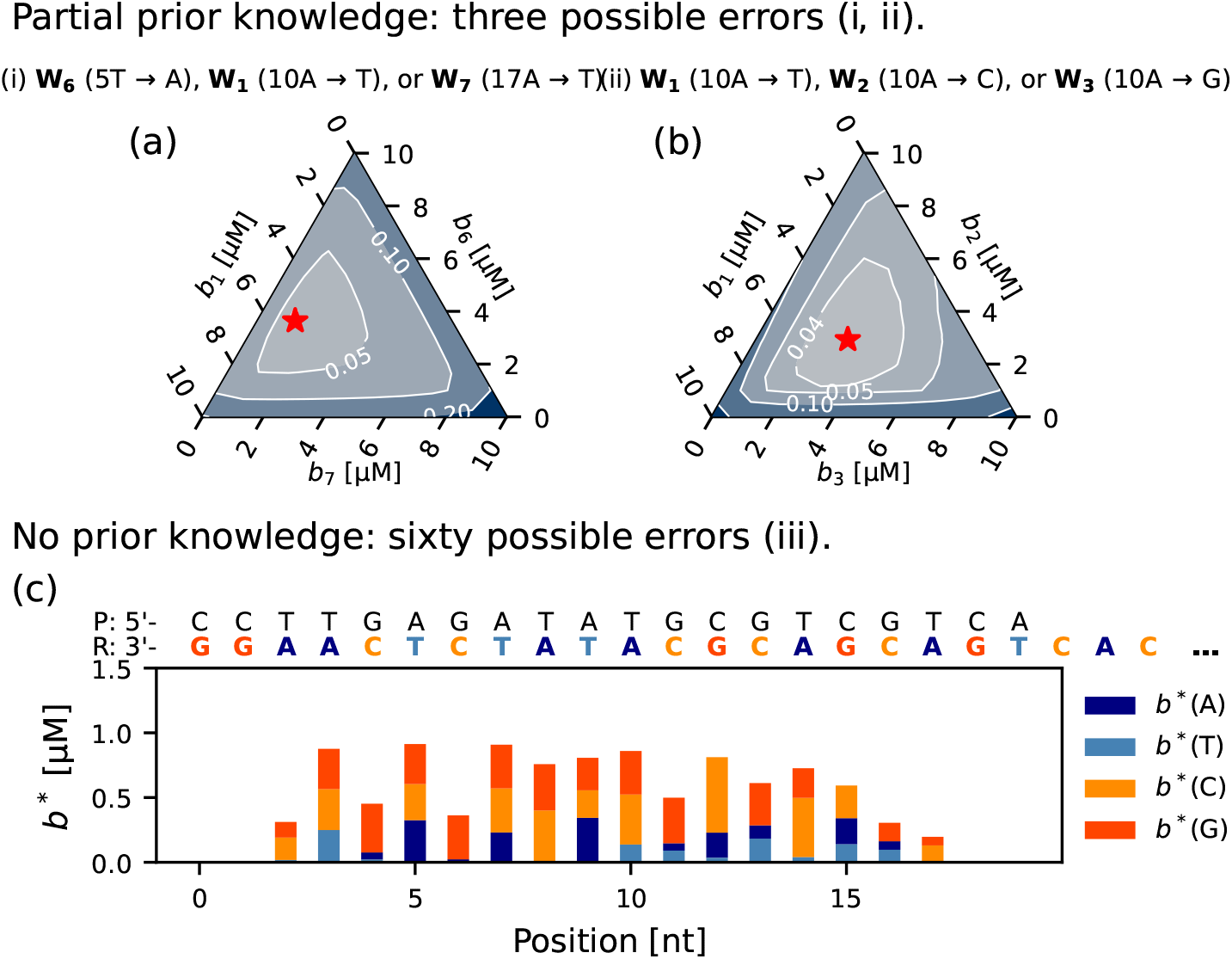
Optimal blocker concentrations minimizing *ε*_mean_ given by (6) under the constraint *b*_tot_ = 10 µM. See Fig. S4 for the optimal blocker concentrations minimizing *ε*_max_. (a), (b) There are possible three mutations at different positions (a) or same position (b). The values of *ε*_mean_ are indicated by colors normalized by the maximum error rate in each plot. The optimal blocker concentrations are indicated by stars. (c) Optimal blocker concentrations for the case with 60 possible errors. The sequences of R and P are shown on the top for the reference. The expressions *b*^∗^(N) (N is A, T, C, or G) indicate the optimal concentrations of the blockers targeting N mutation at each position. *ε*_mean_ without blockers are 0.36 (a), 0.48 (b), and 0.33 (c). The minimum *ε*_mean_ at the optimal blocker concentrations are 0.039 (a), 0.035 (b), and 0.18 (c).

## 4 Optimal blocker combinations

As seen in Fig. 6, *ε*_mean_ and *ε*_max_ have a gradient, suggesting the possibility of minimizing errors by optimizing blocker concentrations. We theoretically derive the optimal blocker distribution 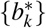 that minimizes the mean or maximum error fraction based on Eq. (6) (see Methods). We obtain 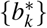 numerically under the constraints *b*_tot_ = ∑ _*j*_ *b* _*j*_ = 10 µM and *b* _*j*_ ≥ 0 for ∀ *j*. It can be shown that *ε*_mean_({*b*_*k*_}) and *ε*_max_({*b*_*k*_}) are convex functions of {*b*_*k*_} for arbitrary 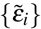 and {*K*_*i,j*_}. This implies that we can efficiently find 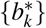 by a gradient descending method, without risking to be stuck in a local minimum. Actually, the *ε*_mean_ and *ε*_max_ have single minima (Fig. 7a, b).

We apply this approach to three situations with different mismatch patterns (Fig. 7 and S4). In addition to the scenarios (i) and (ii) introduced above, where we have partial prior knowledge of the errors, we tested the scenario with no prior knowledge (iii). In all examples, we considered the worst case, where the contamination probability is uniform, *p*_*i*_ = 1*/N*. The results are summarized in Table 1.

In (i) and (ii), the model calculation shows significant error reduction by the optimization (Table 1), implying the effectiveness of the optimization. The obtained optimal blocker combinations are indicated by a star in Fig. 6. The estimated reduction ratio in the experiments was summarized in Table 1. The improvement by the optimization is estimated to be limited. The major reason for this is that the uniform blocker concentrations are already very effective. The addition of blockers with concentrations larger than 3 µM can effectively suppress errors (Fig. 2b–d). We used *b*_tot_ = 10 µM, corresponding to 3.3 µM for each blocker in (i) and (ii) for the uniform blocker concentrations. Thus, we can add a sufficient amount of blockers in the reaction buffer without affecting the amplification of the target sequences. In conclusion, an advantage of the method is that we can significantly suppress errors even without fine optimization of the blocker concentrations. Nevertheless, if an application of the method requires it, the theory we introduced permits to finely optimize error suppression.

Finally, in order to see the limitation of the blocker method, we estimated the error suppression with a mixture of all sixty blocker sequences, assuming that there are sixty error possibilities (Fig. 7c and S4c). We found that the blocker method is still effective, and the optimal blocker combination reduced *ε*_mean_ and *ε*_max_ with reduction ratios of 1.8 and 1.4, respectively (Table 1). Although this scenario (iii) may not be relevant for practical virus testing, it illustrates that the blocker addition always reduces errors even in this worst-case scenario.

The optimal blocker distribution 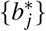 (Fig. 7c) can be explained by the dependence of 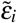 on the position and the dependence of *K*_*i,j*_ on the sequence combinations. In particular, 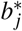 are small at the edges because 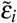 are smaller at the edges than at the internal positions (as shown in Fig. 4). Therefore, the priority to target the edges with blockers is low.

The concentrations 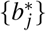 optimizing *ε*_mean_ are strongly biased toward targeting C and G mutations (Fig. 7c). Since C and G bind more strongly than A and T, corresponding *K*_*i,j*_ values are typically small. Therefore, targeting C and G mutations is more effective in reducing *ε*_mean_ under the constraint of a fixed *b*_tot_. This tendency is quantified by the blocking efficacy *α*_*j*_ of the blockers, defined as *α*_*j*_ = ∑_*i*_(1*/K*_*ij*_) (Fig. S5a). 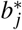 for *ε*_mean_ increases with *α*_*j*_ at small *α*_*j*_. The optimal concentrations 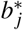 for *ε*_mean_ decrease at large *α*_*j*_ because a small amount is enough for the blockers with large *α*_*j*_ to suppress errors.

In contrast, 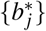 for *ε*_max_ is strongly biased toward targeting A and T mutations (Fig. S4c). Accordingly, 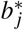 for *ε*_max_ decreases monotonically with *α*_*j*_ (Fig. S5b). In particular, 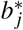 is roughly proportional to 1*/α*_*j*_ (Fig. S5b, inset). This tendency is consistent with intuition as well. If *ε*_*i*_ of a particular template W_*i*_ is larger than other templates, we need to increase *b*_*i*_ to reduce *ε*_max_. The error *ε*_*i*_ is large when B_*i*_ is not effective and, therefore, has a small *α*_*i*_ value. Hence, 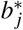 for the blockers with small *α*_*j*_ should be large to minimize *ε*_max_, which results in a uniform *ε*_*i*_ distribution.

## 5 Discussion

We demonstrated the effectiveness of the blocker method in PCR with limited information about the contaminating sequences. Our methodology is based on adding combinations of multiple blocker sequences informed by biophysical considerations. In particular, we successfully modeled the effect of the blockers and obtained the relevant parameters. With this approach, we were able to reduce the mean and maximum error fractions even in the worst-case scenario where all sixty possible contaminations are present (Table 1).

Adding blockers does not increase the error rate unless sufficiently large concentrations of blocker sequences inhibit the replication of R (Fig. 2a). This means that even though the perfect optimization achieves the minimum error fraction, an inaccurate optimization is still effective. In fact, the average error *ε*_mean_ is also reduced by uniform blocker concentrations *b*_*i*_ = *b*_tot_*/N* (Table 1). This tolerance is an inherent advantage of the blocker method because we can reduce the experimental cost at the expense of reducing the error suppression effect. For example, we neglected the dependence of 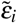 on the nucleotide species and considered only the mismatch position (Fig. 4). The *K*_*i,j*_ values are estimated from the sequences by the nearest neighbor method with a calibration (Fig. 3). While these approximations may reduce the error suppression effect, they significantly lower the burden of evaluating parameters, 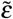 and *K*_*i,j*_. Thus, in applications, assuming that the model of 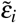 and calibration curve of *K*_*i,j*_ obtained in this work are applicable to other primer sequences, one does not need additional calibration for each system. If necessary, one can further reduce errors by evaluating *K*_*i,j*_ and 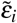 values in each system.

Optimization based on biophysical modeling has not been explored so far in genetic technologies and is itself of conceptual interest. However, for some applications, optimization could bring significant benefits. As mentioned above, the blocker method may be effective in thermally sensitive systems such as genome editing^18^ and RNA interference^19^ in biological cells, where temperature tuning for increasing hybridization specificity is limited. The blocker method targets nucleic acid hybridization and could apply to these gene technologies. However, the maximum blocker concentration in cells may be restricted. In this case, optimization of the blocker concentrations may be essential to suppress errors.

## 6 Methods

The experiments are performed using the same procedure as that in a previous work^10^. The program code (Python) used for finding the optimal blocker concentrations is available on https://github.com/stoyabe/PCR_Blocker.

The experiments consist of two PCR steps (Fig. S1). In the first PCR step, R and W are amplified by linear PCR. In the second PCR step, the amount of the product in the first step, 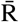 and 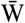, is evaluated by a quantitative PCR (qPCR).

### Linear PCR experiment

In linear PCR, the product amount is amplified linearly because a single primer sequence is used, which simplifies the error analysis (SI Text S1).

DNA strands were synthesized by Eurofins Genomics (sequences are listed in SI Table S1). DNA/LNA chimeric strands were synthesized by Aji Bio-Pharma. The reaction mixture for polymerization contained 25 units*/*ml Hot-start Taq DNA polymerase (New England Biolabs), Taq standard reaction buffer, 2.5 nM R, 2.5 nM W, 100 nM P, and indicated amounts of blocker strands. We performed initial heating for 30 s at 95 °C and, then, 10 cycles of 15 s at 95 °C, 30 s at 60 °C, and 5 s at 68 °C using a PCR cycler (BioRad). Immediately after the cycles, the mixture was cooled down on the ice to stop the enzyme reaction and diluted for quantification.

### Quantification

The second quantitative PCR (qPCR) step was performed on a real-time PCR cycler (BioRad) after the linear PCR experiment for quantifying 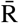 and 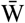 (see SI Text S2). The reaction mixture contains Luna Universal qPCR Master Mix (New England Biolabs), 200 nM each of the primers, and the diluted sample. The dilution rate of the sample was 1/250 in the final concentration. The thermal cycle consisted of initial heating for 60 s at 95 °C, 40 cycles of 15 s at 95 °C, 30 s at 66 °C, and 5 s at 72 °C.

### Optimization of the blocker concentrations

We want to minimize *ε*_mean_ and *ε*_max_ given by Eq. (6) under the constraint that *b*_*i*_≥ 0 for∀ *i* and ∑ _*j*_ *b* _*j*_ = *b*_tot_. It is difficult to obtain the analytical solution under the constraints given by inequalities. Instead, we numerically find the optimum by a gradient descent method. For validating the method’s effectiveness, we verify that *ε*_mean_ and *ε*_max_ are convex functions of {*b*_*i*_}.

#### (i) *Minimization of ε*_mean_

To minimize *ε*_mean_, we first note that the domain{ *b*_*k*_≥ 0∀ *k*; ∑ _*j*_ *b* _*j*_ = *b*_tot_ } is convex, that is, a point between any two points in the domain lies in the same domain. Moreover, the Hessian matrix given by

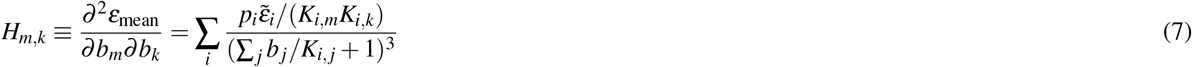

is of the form *H* = *K*^T^*DK*, where *D* is a diagonal matrix with positive entries. Here, *K* is a matrix with the entries *K*_*i,m*_. This implies that the Hessian matrix is nonnegative, hence the function *f* is convex.

The fact that our minimization problem concerns a convex function in a convex domain ensures that the minimum is unique. We can, therefore, find this minimum very efficiently numerically using a gradient descent method. In particular, calling 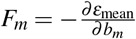, the optimal solution is the unique stable fixed point of the replicator dynamics:

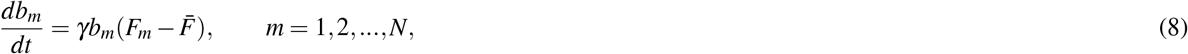

where 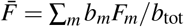 is the average gradient. The coefficient *γ* determines the update rate of {*b*_*m*_}. The second term in the right-hand side of Eq. (8) ensures that the normalization condition ∑ _*j*_ *b* _*j*_ = *b*_tot_ is preserved by the dynamics.

Substituting the explicit expression of the gradient into Eq. (8), we obtain

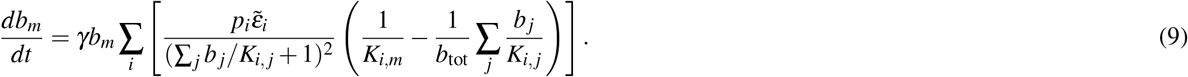

We used the replicator equation (9) to obtain the optimal blocker concentrations that minimize the average error fraction *ε*_mean_.

#### (ii) *Minimization of ε*_max_

The minimization of *ε*_max_ (Eq. (6)) is also possible in a similar manner to *ε*_mean_. We note that *ε*_max_({*b*_*k*_}) is a convex function as well since the maximum of convex functions is a convex function. Let *i*^∗^ be the index with the maximum *ε*_*i*_. Then, the replicator equation reads

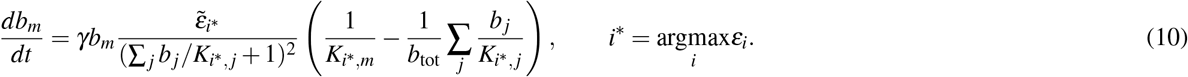

We note that, in the numerical integration of Eq. (10), one has to dynamically update *i*^∗^, i.e., compute at each time step the value of *i* that maximizes *ε*_*i*_.

## Author contributions statement

TT did the experiments. TT, HA, SP, and ST developed the theory, designed the system, and wrote the paper.

## Acknowledgments

ST is supported by JSPS KAKENHI Grant Number JP23K17658 and JST ERATO Grant Number JPMJER2302, Japan. SP is supported by KAKENHI Grant Number JP23H01146.

## Data availability

Data are available from the authors upon reasonable request. The program code (Python) used for finding the optimal blocker concentrations is available on https://github.com/stoyabe/PCR_Blocker.

## References

1. Mullis, K. B. & Faloona, F. A. Specific synthesis of DNA in vitro via a polymerase-catalyzed chain reaction. In Meth. Enzym., 335–350, DOI: 10.1016/0076-6879(87)55023-6 (Elsevier, 1987).

2. Whiley, D. M. & Sloots, T. P. Sequence variation in primer targets affects the accuracy of viral quantitative PCR. J. Clin. Virol. 34, 104–107, DOI: 10.1016/j.jcv.2005.02.010 (2005).

3. Ørum, H. et al. Single base pair mutation analysis by PNA directed PCR clamping. Nucleic Acids Res. 21, 5332–5336, DOI: 10.1093/nar/21.23.5332 (1993).

4. Li, M. et al. Nucleic acid tests for clinical translation. Chem. Rev 121, 10469–10558, DOI: 10.1021/acs.chemrev.1c00241 (2021).

5. Yang, H. et al. A tailored LNA clamping design principle: Efficient, economized, specific and ultrasensitive for the detection of point mutations. Biotech. J. 16, 2100233, DOI: 10.1002/biot.202100233 (2021).

6. Homma, C. et al. Effectiveness of blocking primers and a peptide nucleic acid (PNA) clamp for 18s metabarcoding dietary analysis of herbivorous fish. PLoS one 17, e0266268, DOI: 10.1371/journal.pone.0266268 (2022).

7. Petersen, M. & Wengel, J. LNA: a versatile tool for therapeutics and genomics. Trends Biotech. 21, 74–81, DOI: 10.1016/s0167-7799(02)00038-0 (2003).

8. Pellestor, F. & Paulasova, P. The peptide nucleic acids (PNAs), powerful tools for molecular genetics and cytogenetics. Eur. J. Hum. Genet. 12, 694–700, DOI: 10.1038/sj.ejhg.5201226 (2004).

9. Owczarzy, R., You, Y., Groth, C. L. & Tataurov, A. V. Stability and mismatch discrimination of locked nucleic acid–DNA duplexes. Biochem. 50, 9352–9367, DOI: 10.1021/bi200904e (2011).

10. Aoyanagi, H., Pigolotti, S., Ono, S. & Toyabe, S. Error-suppression mechanism of PCR by blocker strands. Biophys. J. 122, 1334–1341, DOI: 10.1016/j.bpj.2023.02.028 (2023).

11. Hopfield, J. J. Kinetic proofreading: A new mechanism for reducing errors in biosynthetic processes requiring high specificity. Proc. Nat. Acad. Sci. 71, 4135–4139, DOI: 10.1073/pnas.71.10.4135 (1974).

12. Allawi, H. T. & SantaLucia, J. Thermodynamics and NMR of internal G T mismatches in DNA. Biochem. 36, 10581–10594, DOI: 10.1021/bi962590c (1997).

13. SantaLucia, J. A unified view of polymer, dumbbell, and oligonucleotide DNA nearest-neighbor thermodynamics. Proc. Nat. Acad. Sci. 95, 1460–1465, DOI: 10.1073/pnas.95.4.1460 (1998).

14. Cock, P. J. et al. Biopython: freely available Python tools for computational molecular biology and bioinformatics. Bioinformatics 25, 1422–1423, DOI: 10.1093/bioinformatics/btp163 (2009).

15. Kwok, S. et al. Effects of primer-template mismatches on the polymerase chain reaction: Human immunodeficiency virus type 1 model studies. Nuc. Acid. Res. 18, 999–1005, DOI: 10.1093/nar/18.4.999 (1990).

16. Huang, M.-M., Arnheim, N. & Goodman, M. F. Extension of base mispairs by Taq DNA polymerase: implications for single nucleotide discrimination in PCR. Nuc. Acid. Res. 20, 4567–4573, DOI: 10.1093/nar/20.17.4567 (1992).

17. Bru, D., Martin-Laurent, F. & Philippot, L. Quantification of the detrimental effect of a single primer-template mismatch by real-time PCR using the 16s rRNA gene as an example. Appl. Environ. Microbiol. 74, 1660–1663, DOI: 10.1128/aem.02403-07 (2008).

18. Anzalone, A. V., Koblan, L. W. & Liu, D. R. Genome editing with CRISPR-Cas nucleases, base editors, transposases and prime editors. Nat. Biotech. 38, 824–844, DOI: 10.1038/s41587-020-0561-9 (2020).

19. Agrawal, N. et al. RNA interference: Biology, mechanism, and applications. Microbiol. Mol. Biol. Rev. 67, 657–685, DOI: 10.1128/mmbr.67.4.657-685.2003 (2003).

