## Supplementary methods and figures. for "Blocking uncertain mispriming errors of PCR"

### S1 Supplementary Methods: Linear PCR

Figure S1 shows the flow of the experiments.

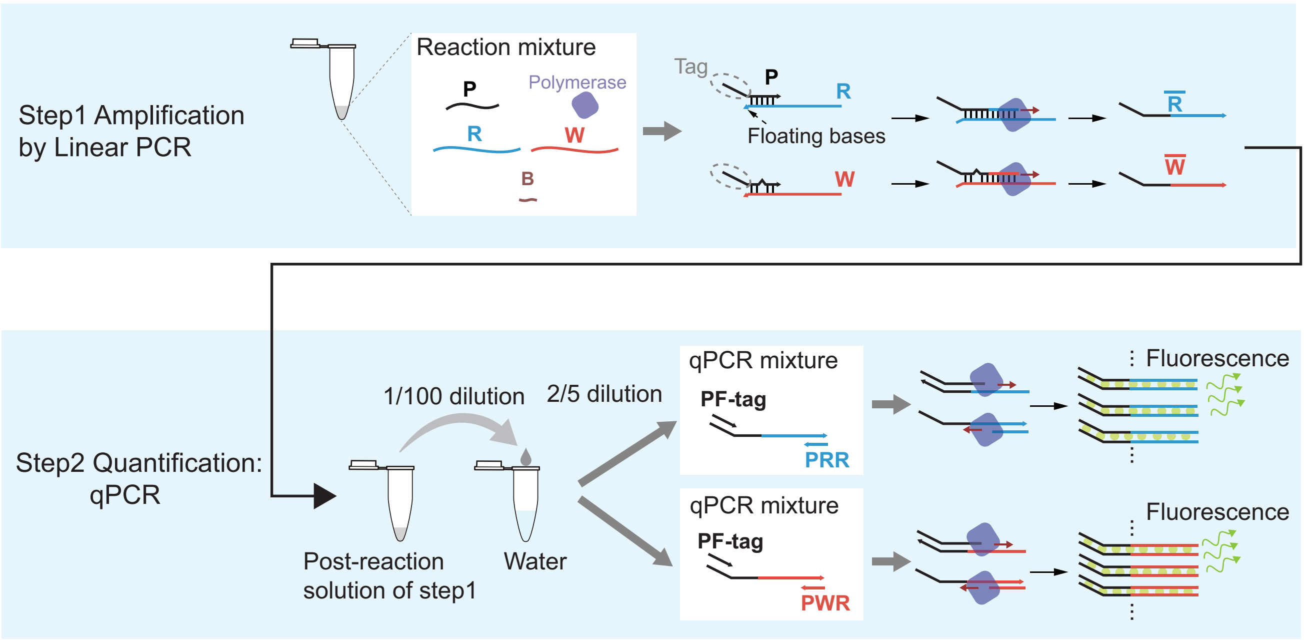


**Figure S1.** Scheme of the experiments. R and W have similar sequences in their primer region and are otherwise different. P includes a tag sequence for the subsequent quantitative PCR (qPCR). Two bases at the 3’ ends of R and W are floating to prevent elongation of the primer tag sequence.

#### Sequences for the linear PCR

Sequences of the DNA strands used in the linear PCR step (Fig. S1) are listed in Table S1. The sequences of the primer-binding regions of R and W differ by a single base, resulting in a single-base mismatch between P and W. The sequences outside the primer-binding region of R and W are different. The primer P includes a tag sequence at its 5’ end for the subsequent quantification step. The polymerase elongates the primers of the hybridized complexes P:R and P:W to produce $\overline{\mathrm{R}}$ and $\overline{\mathrm{W}}$, respectively. The 3’ ends of R and W possess two-base overhang to prevent elongation from these ends.

The blockers are LNA-DNA chimeric strands and hybridize to a part of the primer-binding regions of R and W. The addition of LNA nucleotides increases the specificity of the hybridization. The 3’-ends have two-base floating sequences that are not complementary to the template sequences, so the blockers do not work as primers.

**Table S1.** The DNA sequences used in the linear PCR step. Mutations in W are indicated by bold characters. LNA bases are indicated by lowercase.

| Name | Mutation | Sequence |
|---|---|---|
| R | | 5' – CTCTC GTTCT CAACC TACCA TGGCA AGGGT TGTTC TTTGT TGGTG TCCAC TGACG ACGCA TATCT CAAGG TA – 3' |
| $W_1$ | 10A → T | 5' – TACAT ATGAC GCACA GATGC AGCCA ACTCC ACTCA CTTCT CTCTT CAGAT TGACG ACGC**T** TATCT CAAGG TA – 3' |
| $W_2$ | 10A → C | 5' – ··· TGACG ACGC**C** TATCT CAAGG TA – 3' |
| $W_3$ | 10A → G | 5' – ··· TGACG ACGC**G** TATCT CAAGG TA – 3' |
| $W_4$ | 2A → T | 5' – ··· TGACG ACGCA TATCT CA**T** GG TA – 3' |
| W | 2A → G | 5' – ··· TGACG ACGCA TATCT CA**G** GG TA – 3' |
| W | 2A → C | 5' – ··· TGACG ACGCA TATCT CA**C** GG TA – 3' |
| $W_6$ | 5T → A | 5' – ··· TGACG ACGCA TATC**A** CAAGG TA – 3' |
| W | 5T → G | 5' – ··· TGACG ACGCA TATC**G** CAAGG TA – 3' |
| W | 5T → C | 5' – ··· TGACG ACGCA TATC**C** CAAGG TA – 3' |
| W | 14A → T | 5' – ··· TGACG **T**CGCA TATCT CAAGG TA – 3' |
| W | 14A → G | 5' – ··· TGACG **G**CGCA TATCT CAAGG TA – 3' |
| W | 14A → C | 5' – ··· TGACG**C**CGCA TATCT CAAGG TA – 3' |
| $W_7$ | 17A → T | 5' – ··· TG**T**CG ACGCA TATCT CAAGG TA – 3' |
| W | 17A → G | 5' – ··· TG**G**CG ACGCA TATCT CAAGG TA – 3' |
| W | 17A → C | 5' – ··· TG**C**CG ACGCA TATCT CAAGG TA – 3' |
| $W_5$ | 19T → A | 5' – ··· **A**GACG ACGCA TATCT CAAGG TA – 3' |
| W | 19T → G | 5' – ··· **G**GACG ACGCA TATCT CAAGG TA – 3' |
| W | 19T → C | 5' – ··· **C**GACG ACGCA TATCT CAAGG TA – 3' |
| $\overline{R}$ | | 5' – GTCAC CCAAT CGTCC TAGAT CCTTG AGATA TGCGT CGTCA GTGGA CACCA ACAAA GAACA ACCCT TGCCA TGGTA GGTTG AGAAC GAGAG – 3' |
| $\overline{W}_1$ | | 5' – GTCAC CCAAT CGTCC TAGAT CCTTG AGATA TGCGT CGTCA ATCTG AAGAG AGAAG TGAGT GGAGT TGGCT GCATC TGTGC GTCAT ATGTA – 3' |
| P | | 5' – GTCAC CCAAT CGTCC TAGAT CCTTG AGATA TGCGT CGTCA – 3' |
| $B_1$ | | 5' – GAGAT a**a**gcG TCGTT T – 3' |
| $B_2$ | | 5' – GAGAT a**g**gcG TCGTT T – 3' |
| $B_3$ | | 5' – GAGAT a**c**gcG TCGTT T – 3' |
| $B_4$ | | 5' – Cc**a**tg AGATA TGCGA T – 3' |
| $B_5$ | | 5' – GTCGT c**t**atC TGAAT T – 3' |
| $B_6$ | | 5' – CCTTg **t**gaTA TGCGA T – 3' |
| $B_7$ | | 5' – TGCGT Cg**a**ca ATCTT T – 3' |

### S2 Supplementary Methods: Quantitative PCR

#### Sequences

We use two primer sets {PF-tag, PRR} and {PF-tag, PWR} (Fig. S1, Table S2) for detecting $\overline{\mathrm{R}}$ and $\overline{\mathrm{W}}$, respectively. The PRR and PWR hybridize to the 21-base region at the 3' ends of $\overline{\mathrm{R}}$ and $\overline{\mathrm{W}}$, respectively. The reaction mixture contains diluted sample of the linear PCR (diluted to 1/250 in the final concentration), 200-nM each primers and Luna Universal qPCR Master Mix (New England Biolabs).

**Table S2.** The DNA sequences for the quantitative PCR.

| Name | Sequence |
|---|---|
| PF-tag | 5' – GTCAC CCAAT CGTCC TAGAT C – 3' |
| PRR | 5' – CTCTC GTTCT CAACC TACCA T – 3' |
| PWR | 5' – TACAT ATGAC GCACA GATGC A – 3' |

#### Standard curves

We measured the standard curves used for the quantification of $\overline{\mathrm{R}}$ and $\overline{\mathrm{W}}$ (Fig. S2). P, R, and W used in the linear PCR step are carried over to the quantitative PCR step. Although they are diluted to 1/250, they increase the background level and affect the quantification when the template concentration is low. Hence, for making the standard curves, we added 400 pM P and 6.25 pM R and 6.25 pM W to imitate the effect of the carry over. In addition to these strands, the reaction mixture contains Luna Universal qPCR Master Mix (New England Biolabs), templates $\overline{\mathrm{R}}$ or $\overline{\mathrm{W}}$, and 200-nM each primer sets, {PF-tag, PRR} and {PF-tag, PWR}. The template concentrations were 160, 20, 2.5, 0.31, 0.039, and 0.049 pM.

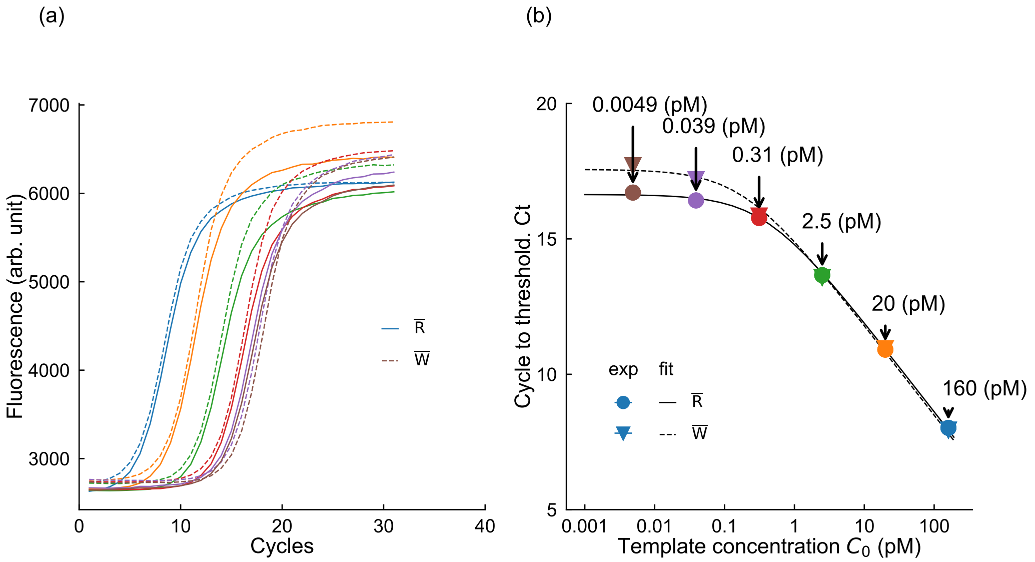


**Figure S2.** Standard curve measurement. (a) The qPCR curves with different initial template concentrations. (b) Standard curves. $\mathrm{C_t}$ values saturate at low template concentrations due to mainly the contamination of the residues of the linear PCR step. We directly fitted these points with Eq. (S1) with $\delta$ and $\phi$ as the fitting parameters (solid curves).

The $\mathrm{C_t}$ values, which are defined by the cycle number where the fluorescence reached a threshold value, are related to the template concentration $C_0$ by

$$\mathrm{C_t} = -\log_2[C_0 + \delta] + \phi. \tag{S1}$$

We calculated $\mathrm{C_t}$ based on the maxRatio method[1]. We fitted the experimental data by this relation (Fig. S2b) and obtained the background level $\delta$ (15.3 for R and 15.1 for W) and intercept $\phi$ (0.386 for R and 0.186 for W). Using these parameters, $[\overline{\mathrm{R}}]$ and

$[\overline{\mathrm{W}}]$ are calculated from $\mathrm{C_t}$ values by

$$[\overline{\mathrm{R}}] = 2^{-(\mathrm{C_t}-15.3)} - 0.386,$$
$$[\overline{\mathrm{W}}] = 2^{-(\mathrm{C_t}-15.1)} - 0.186. \tag{S2}$$

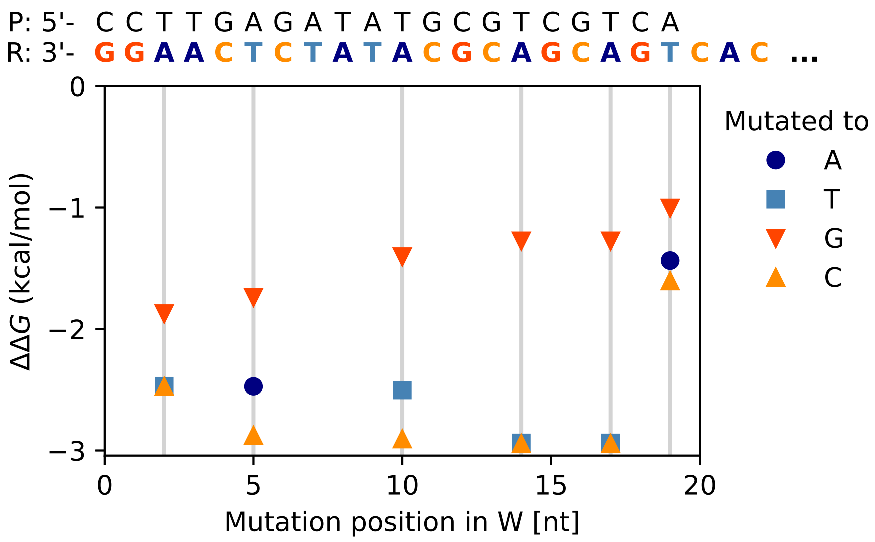


**Figure S3.** The free energy difference $\Delta\Delta G = \Delta G(\mathrm{P}:\mathrm{R})v - \Delta G(\mathrm{P}:\mathrm{W}_i)$ for each mutation calculated by the nearest neighbor method[2,3]. Smaller $\Delta\Delta G$ values mean higher stability of the mismatched hybridization.

### S3 Supplementary Figures

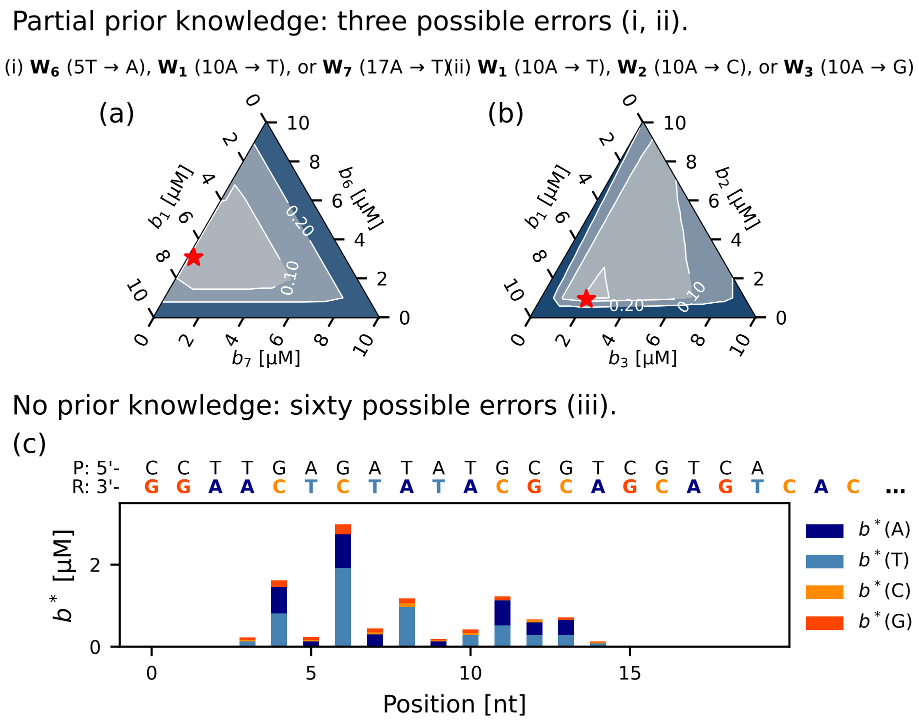


**Figure S4.** Same plot as Fig. 6 but for $\varepsilon_{\mathrm{max}}$. The minimum error fractions with the optimal blocker concentrations are 0.048 (a), 0.044 (b), and 0.34 (c). The error fractions in the absence of the blockers are 0.48 (a), 0.48 (b), and 0.48 (c).

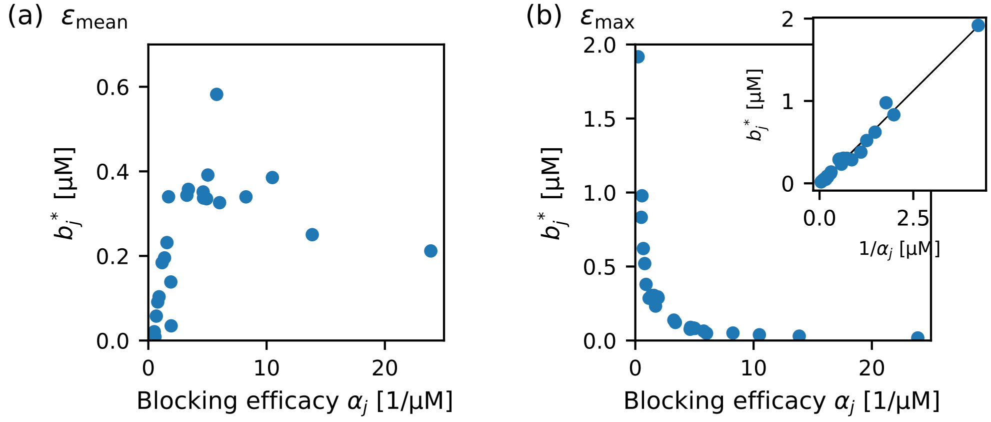


**Figure S5.** Optimal blocker concentrations for $\varepsilon_{\mathrm{mean}}$ (a) and $\varepsilon_{\mathrm{max}}$ (b) in the situation (iii) are plotted against the blocking efficacy $\alpha_j = \sum_i (1/K_{ij})$. The data is limited to the blockers targeting the intermediate regions (from 7-th to 14-th nucleotides from the 3'-end) because edge nucleotides are affected by $\tilde{\varepsilon}_i$. The solid curve in the inset of (b) is a linear fit.

### References


1. Shain, E. B. & Clemens, J. M. A new method for robust quantitative and qualitative analysis of real-time PCR. *Nuc. Acid. Res.* **36**, e91–e91, DOI: 10.1093/nar/gkn408 (2008).

**2.** SantaLucia, J. A unified view of polymer, dumbbell, and oligonucleotide DNA nearest-neighbor thermodynamics. *Proc. Nat. Acad. Sci.* **95**, 1460–1465, DOI: 10.1073/pnas.95.4.1460 (1998).

**3.** Allawi, H. T. & SantaLucia, J. Thermodynamics and NMR of internal G·T mismatches in DNA. *Biochem.* **36**, 10581–10594, DOI: 10.1021/bi962590c (1997).
